# AVITI library prep miniaturization and combining with Illumina data for phylogenomic and population genomic analyses

**DOI:** 10.1101/2025.07.30.667782

**Authors:** Jacob B. Landis, Ella Hufnagel, Josh M. Felton, Julianna J. Harden, Deivid Almeida, Chelsea D. Specht

## Abstract

Recent advancements in next generation sequencing approaches allow for expansion of evolutionary research into the discovery of genetic patterns and processes underlying diversification across scales. The increased popularity of the Element Bioscience AVITI platform, partially due to the high sequencing accuracy and low cost of reagents, is becoming a viable alternative approach for generating massive amounts of comparative sequencing data across diverse organismal lineages. Using a data set of five accessions from the monocot genus *Costus*, we tested miniaturization conditions for generating robust, cost-effective libraries and made comparisons of data generated by AVITI and Illumina sequencing platforms to investigate the potential for combining data for population genomic and phylogenomic analyses. Our results show that the AVITI and Illumina data sets are highly congruent in terms of inferring overlapping SNPs, with only a small fraction picked up by only one of the two platforms. The rates of duplication in miniaturized libraries were much higher than in full volume libraries and in the Illumina libraries, resulting in missing SNPs and less sequence coverage when volumes are reduced. For all generated libraries, most downstream evolutionary analyses, including clustering algorithms (such as PCA) and phylogenetic inference, yielded similar results. However, Structure analyses were less consistent across datasets, with data from the most miniaturized libraries being assigned to the wrong clusters. The AVITI platform should be seen as a cost-effective approach for generating genomic data for comparison across taxonomic lineages, even for ongoing projects where Illumina data already exists.

## Introduction

Advancements and optimization of next generation sequencing (NGS) technology have allowed a broad expansion of the types of evolutionary questions that can be addressed with sequencing data, in particular the scale of comparative genomic projects that can be completed on limited budgets. The vast majority, in fact over 90%, of all generated NGS data to date have been generated on one of several available Illumina platforms, with MGI-BGISEQ 500 and Element Biosciences AVITI becoming more widely used recently [1]. The Element AVITI, which uses avidity sequencing instead of sequencing by synthesis, is becoming a feasible alternative to Illumina sequencing, requiring less reagent consumption while producing high quality base calls with an accuracy ranging between 95.7 and 97% (one error per 1,000 base pairs) [2,3]. Recent comparisons indicate that AVITI achieves greater variant calling accuracy when compared to Illumina, especially at a depth of 20-30x coverage, with drastic improvements for tandem repeats and homopolymers [4]. Studies comparing genotype calls between AVITI and Illumina GBS sequencing runs show high concordance, with a 91.1% overlap in the number of retained SNPs, with the largest differences between the two platforms associated with the handling of indels [5]. While several studies have shown a high degree of overlap in generated reads and genotype calls, these studies have not tested the downstream implications of using a mixed data set (one that includes both AVITI and Illumina generated reads) for either population genomic or phylogenomic studies.

Combining data sets from different sequencing approaches, typically generated by different library preparation methods, has been shown to produce mixed results. Combining genome resequencing data with target capture data has worked for large scale phylogenomic analyses such as resolving the angiosperm tree of life [6]. Combining whole genome resequencing data with GBS data has been shown to work in cases where the data sets are predominately made up of one type of data, such as adding a handful of resequencing samples to a GBS data set in *Glycine* [7]. However, in cases where the sequencing data are more evenly split between the two sequencing approaches, the robustness of the results can be mixed. Friel et al. [8] found strong signal supporting clustering of sequences of the same specimen generated by different methods (i.e., sister relationships in a phylogenetic tree between GBS and resequencing of the same sample), but found low support in resolving relationships between specimens (i.e., deeper nodes in the phylogenetic tree). Alternatively, as presented at the Botany 2024 conference by Landis and colleagues (unpublished), when analyzing mixed data sets, sequences clustered predominately by library preparation method (GBS v. resequencing) rather than revealing biologically meaningful relationships among sequenced accessions. This relationship was recovered regardless of how the SNPs were called, i.e., whether using a reference-based approach or a reference-free k-mer approach. With recent advancements in sequencing technology resulting in increases in read depth and accuracy with lower costs, finding ways to combine heterogeneous datasets is going to become increasingly important, especially for long-running projects looking for cheaper alternatives for data generation over time and for researchers interested in add new data to existing published or publicly available data sets [9].

In an attempt to reduce costs and increase the number of accessions used in NGS experiments, one commonly used approach is library miniaturization, or using partial volumes for making libraries [10]. Previous comparisons using Illumina DNA preparation kits have shown that sequence quality is not impacted by library miniaturization and that the data generated from reduced volume reactions are generally compatible with data using full volumes as recommended by the manufacturer [11]. Several studies have utilized miniaturization of Roche KAPA library kits [12,13] for different scales of evolutionary projects including genome wide association and phylogenomic analyses. Comparisons down to 1/8th miniaturization have highlighted a strong concordance in recovered genotypes and genotype imputation across miniaturization levels and commercial kits [14].

To test the potential and impacts of combining newly generated AVITI sequencing data to a pre-existing data set and to test for the effects of miniaturization of library preparation, a study system must have well-established genomic resources to serve as the control. The monocot genus *Costus* (Costaceae) provides such a system, with genomic data existing to address evolutionary questions above and below the species level. The genus contains 76 species [15], with a large Neotropical clade [13] radiating after a single long-distance dispersal [16,17] and a basal African grade comprising 24 species [18]. The phylogenetic relationships of the group have been well-characterized with multiple studies using Hyb-seq approaches of hundreds to thousands of loci [13,19,20]. Multiple genomes for the genus have been published [13,21], as well as high-quality transcriptomes [13,22,23] that can be used for candidate gene discovery and for genome annotation. Using the available genomic resources, genome resequencing for SNP calling has been used to investigate differences between bird and bee pollination [13], as well as to phylogenetically place newly described species Specht et al. Monocots 2024 conference (unpublished).

The goals of this study were two-fold. The first is to determine if sequencing data from the AVITI platform can be combined with previously generated Illumina data for downstream population genomic and phylogenomic analyses to produce biologically meaningful results. The same accessions for five different species were sequenced using both platforms using both freshly collected leaf tissue stored at -80℃ and silica-dried leaf tissue stored at room temperature. The second goal is to test miniaturization of the Element Elevate library prep kit using enzymatic fragmentation from Full down to 1/5th reactions to determine what impacts miniaturization have on generated sequencing reads and downstream analyses, specifically to address at what scale miniaturization may serve as a feasible cost cutting approach.

## Methods

A previous study presented and described the generation of ∼80x Illumina sequencing data for 20 species of *Costus* to investigate macroevolutionary questions associated with pollination syndromes [13]. Given that one AVITI flow cell can generate ∼300 Gbp and the target was 10x coverage per library (each species of *Costus* has approximately a 1 Gbp genome), five taxa (*Costus beckii* R3394, *Costus lima* R3362, *Costus spiralis* R3218, *Costus villosissimus* R3074, and *Costus whiskeycola* R3026) were chosen for comparisons of sequencing data generated by Illumina NovaSeq and AVITI using five different library miniaturization settings. DNA extractions were made from freshly collected leaves stored frozen at -80℃ were used for four species (*C. beckii*, *C. spiralis*, *C. villosissimus*, and *C. whiskeycola*), and from leaves stored in silica gel at room temperature for *C. lima* and *C. spiralis*. DNA from all accessions was extracted in four replicates using a modified SDS protocol [24,25] that has been used previously for phylogenetic studies in *Costus* [13,19]. For each accession, two rounds of extractions were done, with the second round generally showing lower yields than the first [26]. Extracted DNA was quantified using the Qubit double stranded Broad Range (Invitrogen, Waltham, MA, USA). Quality of DNA was determined by running 4 µL on a 1.5% agarose gel at 120 volts for 45 minutes. Replicates for the same accession and the same round of extractions were pooled together and quantified again prior to library construction.

Illumina library preparation and sequencing was previously described by Valderrama et al. [13] for the accessions used in this study. Library prep for AVITI was done using the Element Elevate kit with the Elevate long UDI Adapter Kit Set A. One of the major goals of this study was to investigate the efficiency and applicability of kit miniaturization, with five different levels used: full libraries, 4/5th concentration libraries, 3/5th concentration libraries, 2/5th concentration libraries, and 1/5th concentration libraries. After discussing methodological options with specialists from Element Biosciences, volumes were held constant across all levels by adding the necessary water to reach the final volumes in the standard library kit. Keeping volumes the same and doing more than one library at a time removed the need to pipette very small amounts of liquids multiple times, diminishing pipetting errors. Total input DNA was also scaled with the different levels of library miniaturization spanning 250 ng, 200 ng, 150 ng, 100 ng, and 50 ng total; all in 40 µL of water. In addition to altering input concentrations, the standard Elevate library kit protocol was modified with the following modifications: fragmentation was extended to five minutes at 60℃ to target ∼500 bp (instead of four minutes), working volume of libraries were raised to 100 µL by adding in 50 µL low TE prior to size selection, PCR amplifications were done at half reactions, with 25 µL of water added to the PCR product prior to final bead cleaning, and the final cleaned libraries were resuspended in 17 µL of low TE instead of 20 µL low TE. The number of PCR cycles varied between 8-24 cycles depending on the library concentration (Table 1), with the goal of achieving a minimum of 1-5 ng/µL after the final cleanup. The final libraries were quantified using the Qubit double stranded Broad Range kit and select libraries were run on a Bioanalyzer to check for a proper size distribution.

**Table 1.**
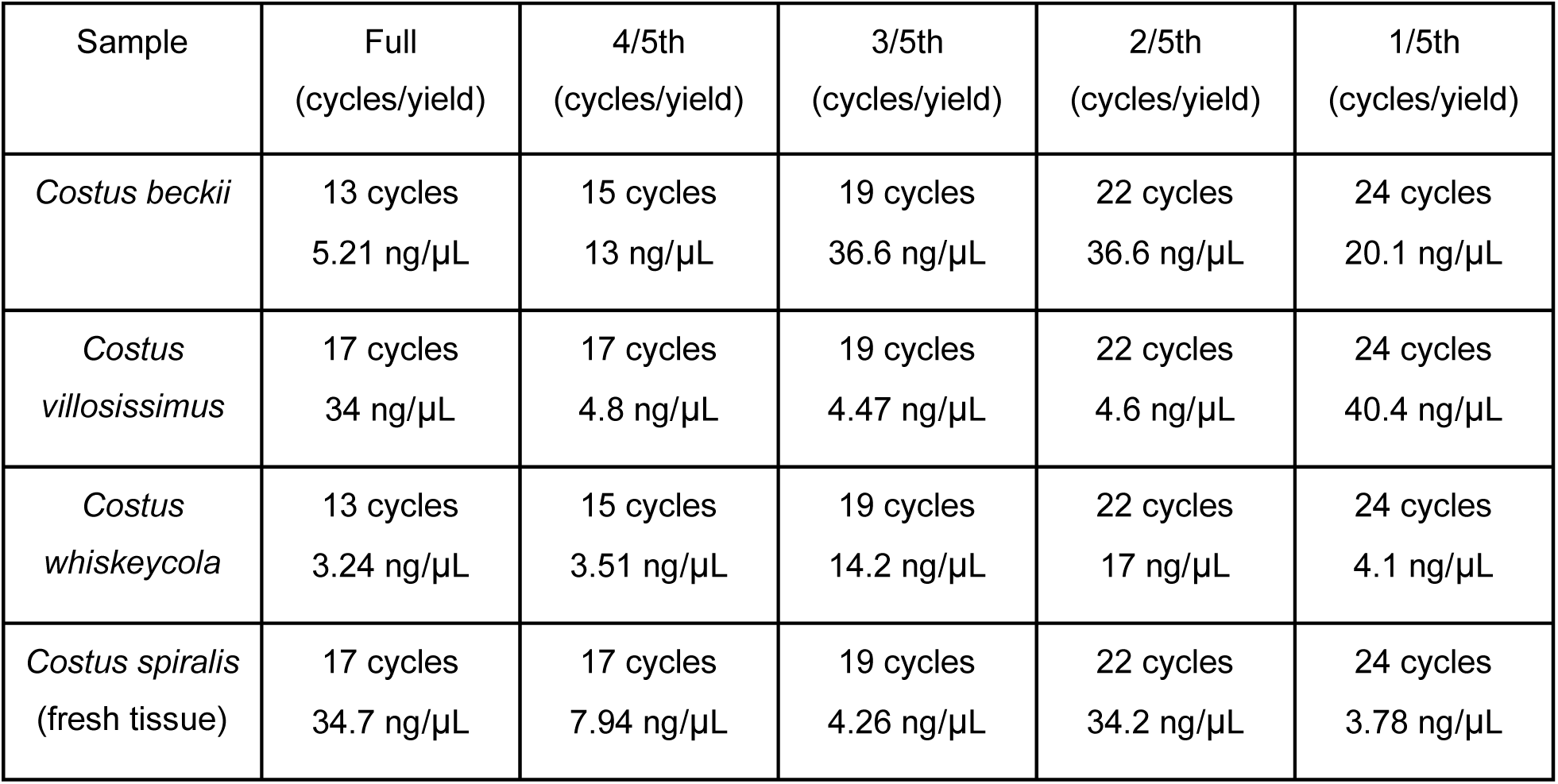
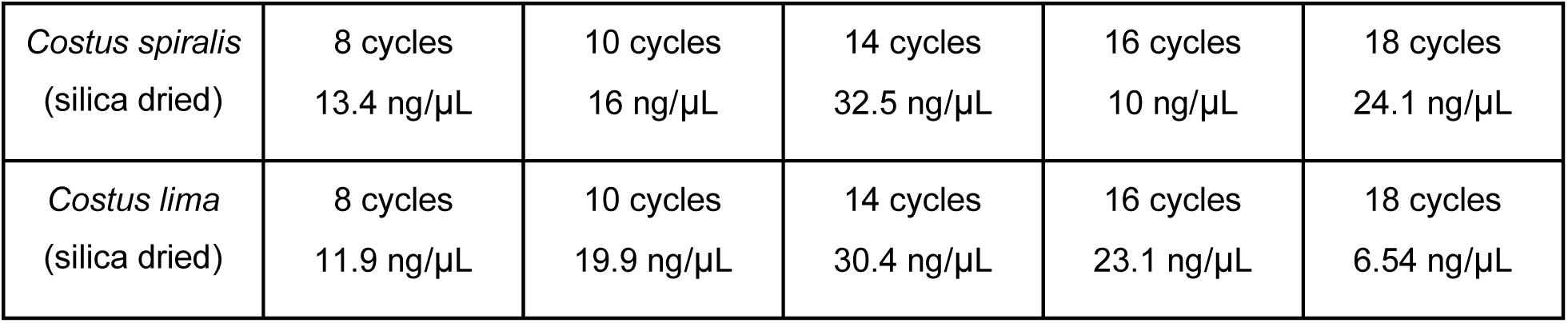
Summary of the number of cycles used for library amplification and the yield (ng/µL) after library preparation for each accession and each library threshold.

| Sample | Full<br>(cycles/yield) | 4/5th<br>(cycles/yield) | 3/5th<br>(cycles/yield) | 2/5th<br>(cycles/yield) | 1/5th<br>(cycles/yield) |
| --- | --- | --- | --- | --- | --- |
| *Costus beckii* | 13 cycles<br>5.21 ng/μL | 15 cycles<br>13 ng/μL | 19 cycles<br>36.6 ng/μL | 22 cycles<br>36.6 ng/μL | 24 cycles<br>20.1 ng/μL |
| *Costus villosissimus* | 17 cycles<br>34 ng/μL | 17 cycles<br>4.8 ng/μL | 19 cycles<br>4.47 ng/μL | 22 cycles<br>4.6 ng/μL | 24 cycles<br>40.4 ng/μL |
| *Costus whiskeycola* | 13 cycles<br>3.24 ng/μL | 15 cycles<br>3.51 ng/μL | 19 cycles<br>14.2 ng/μL | 22 cycles<br>17 ng/μL | 24 cycles<br>4.1 ng/μL |
| *Costus spiralis*<br>(fresh tissue) | 17 cycles<br>34.7 ng/μL | 17 cycles<br>7.94 ng/μL | 19 cycles<br>4.26 ng/μL | 22 cycles<br>34.2 ng/μL | 24 cycles<br>3.78 ng/μL |
| *Costus spiralis*<br>(silica dried) | 8 cycles<br>13.4 ng/μL | 10 cycles<br>16 ng/μL | 14 cycles<br>32.5 ng/μL | 16 cycles<br>10 ng/μL | 18 cycles<br>24.1 ng/μL |
| *Costus lima*<br>(silica dried) | 8 cycles<br>11.9 ng/μL | 10 cycles<br>19.9 ng/μL | 14 cycles<br>30.4 ng/μL | 16 cycles<br>23.1 ng/μL | 18 cycles<br>6.54 ng/μL |

The 30 libraries, five for each miniaturization test for five accessions with one accession consisting of both fresh and silica dried material, were pooled by adding 20 ng total from each library followed by a bead cleanup to reduce volume. The pool was run on an AVITI Freestyle 2x150 flow cell. Raw reads were converted to fastq and demultiplexed using Bases2fastq v2.1.0 (Element Biosciences, San Diego, CA) allowing one mismatch in the reverse index. Demultiplexed reads were cleaned using fastp v0.23.4 [27] with auto detection of adapters, a minimum read length of 75 bp, a minimum quality of 20, and removing any polyG strings. Cleaned reads were mapped to the *C. bracteatus* genome (CoGe ID 63651) [13] using bwa-mem2 v2.2.1 [28], with the resulting mapping file converted to BAM and sorted with Samtools v1.19.2 [29]. Duplicates were marked using the version of piccard (http://broadinstitute.github.io/picard/) packaged with GATK 4.6.0.0 [30]. SNPs were called on the duplicates marked files using the mpileup and call commands in BCFtools v1.19 [31].

VCFtools v0.1.16 [32] was used to filter the 125,131,583 initially called SNPs to only keep high quality, unlinked SNPs. Initial filtering allowed a maximum missing of 20% for any site, keeping only biallelic sites by removing indels, a minimum depth of 3, minimum quality of 20, a minor allele frequency of 0.05, and a minor allele count of three. Remaining SNPs were filtered for linkage disequilibrium using plink 1.9 [33] with a r^2^ of 0.5, a 20 kb window, and a 10 kb step size. The resulting fasta file was converted to fasta and nexus format for downstream analyses using vcf2phyllip [34] and structure format using Stacks 2.66 [35].

To investigate shared and unique SNPs generated for each library, files were filtered individually to remove missing data. After keeping only the rows specifying SNP position by removing header rows with awk, Venn diagrams were created with the R package VennDiagram v1.7.3 [36]. For the limited number of cases that SNPs were retained in the AVITI full libraries compared to their Illumina counterparts (see below), the duplicates marked mapping files were used to manually investigate differences between AVITI and Illumina using the Integrative Genomics Viewer (IGV) v2.19.4 [37–39].

Genome wide heterozygosity was calculated for each sequenced library with angsd v0.940-dirty [40] using the options ‘-dosaf 1 -gl 1 -minMapQ 30 -minQ2 20’ to include only high quality mapping sites. The maximum likelihood site frequency spectrum was calculated using the realSFS function of angsd, and then heterozygosity was calculated as the proportion of heterozygous sites out of the total high quality mapped sites.

### Downstream analyses

Phylogenomic analyses were conducted with raxml-ng v1.2.1 [41] using the genotype model of molecular evolution (GTGTR) and 100 bootstrap replicates. Three separate analyses with different subsets of the data were used to ensure the robustness of the results, all with the 639,116 LD pruned SNP set: the first consisted of the sampling from Valderrama et al. [13] representing 20 species of *Costus* plus the six accessions newly sequenced with AVITI, the second included just the accessions with AVITI sequences and the same-species Illumina sequences rooted with *C. malortieanus*, and the third was formed from the previous dataset but including only sequences generated from Illumina and the full AVITI libraries (i.e. miniaturized AVITI libraries were excluded). The three datasets were analyzed to test for robust phylogenetic placement and to ensure results of clustering analyses were not impacted by the proportion of the data set comprising AVITI sequencing data. In addition to phylogenetic inference, the fasta file of all generated libraries was used to calculate pairwise distances across libraries using Mega11 [42].

A Principal Component Analyses (PCA) was used to identify closest genetic clusters in the R package SNPRelate v1.22.0 [43] with the input being the LD pruned VCF file consisting of the AVITI sequence data for each accession and its taxonomically relevant Illumina counterpart, plus *C. malortieanus* as the outgroup. Three separate plots for each data set were inferred: PC1 vs PC2, PC2 vs PC3, and PC3 vs PC4 using ggplot2 v3.5.2 [44]. In addition to PCA, population structure was analyzed with the R package LEA v3.10.2 [45]. The optimal K was determined using the cross-entropy criterion testing K between 1 to 25. The K value that minimized cross-entropy was chosen as the optimal K. Admixture bar plots were created for K=2 to K=10 to show variation.

## Results

After two rounds of SDS extractions followed by pooling, DNA yields ranged from 3.37 to 13.7 ng/µL from fresh tissue and 10.4 and 23.5 ng/µL from silica-dried tissue. As expected, the fresh tissue produced higher molecular weight DNA with less shearing than the silica-dried tissue (Figure 1). The number of cycles needed for library amplification to achieve ∼5 ng/µL varied between DNA input levels, but also between the fresh tissue and the silica-dried tissue (Table 1). Across the board, the 1/5th libraries required 24 cycles for amplification when using fresh tissue, but only 18 cycles when using silica-dried tissue. Generally, for every increase of 50 ng of input DNA, the number of cycles for library amplification could be reduced by two. Extractions that generated low output or required additional cycles to achieve the desired concentration typically had larger sized fragments than expected (Supplemental Figure S1), indicating suboptimal fragmentation during library preparation.

**Figure 1.**
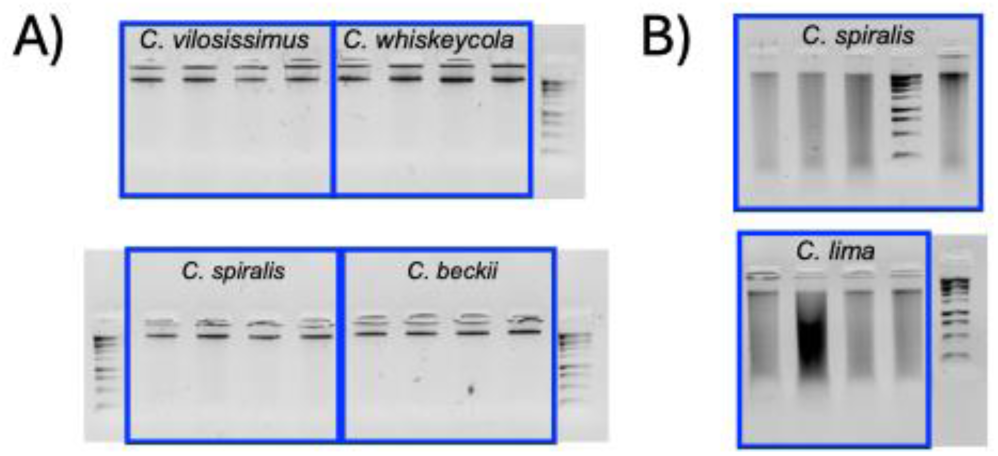
Agarose gel of DNA extractions from fresh (A) and silica-dried (B) tissue with a 1 KB ladder (VWR, Radnor, PA, USA). Box outlines show the accessions used, with the four replicates pooled prior to library preparation. Freshly collected tissue was stored at -80℃, while silica-dried tissue was stored at room temperature.

**Supplemental Figure S1.**
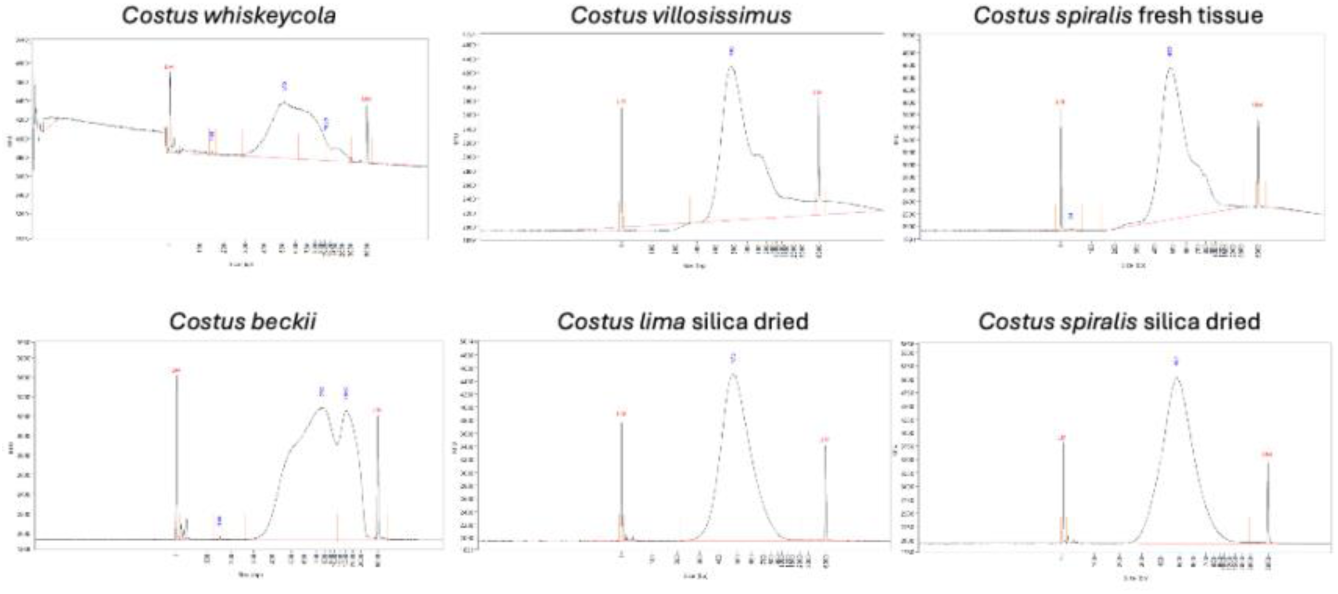
Bioanalyzer traces for the full AVITI libraries for each of the six accessions. *Costus beckii* and *Costus whiskeycola* performed poorly in terms of sequencing output and were also the least uniform in size distribution. The remaining four libraries were all in the expected size distribution of 400-500 bp.

### Sequencing data, SNP calling, and SNP filtering

Pooling the 30 AVITI libraries and running on a Freestyle flow cell 2x150 yielded 312.6 Gbp of total sequencing data. Allowing one mismatch in barcodes resulted in 2.43 to 24.85 Gbp per library with an average of 10.42 Gbp (Figure 2). All libraries were pooled to equal concentrations (20 ng total), but not to equal molar, which contributed to the unequal sequencing output across all libraries.

**Figure 2.**
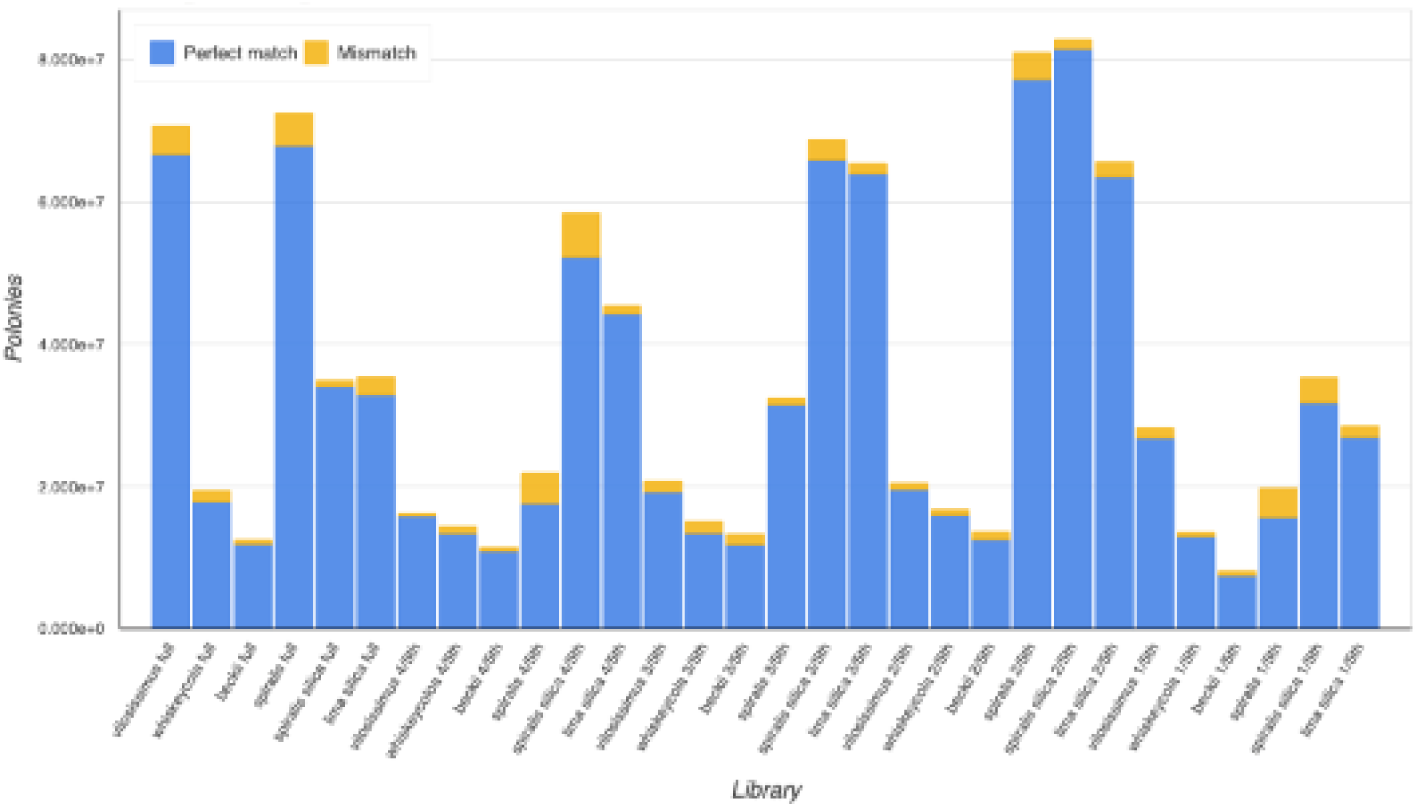
Demultiplexing results from bases2fastq showing the number of polonies per library. Number of polonies x 300 bp gives the approximate yield in Gbp per library with the average being 10.42 Gbp (3.47 million polonies).

After mapping cleaned reads for both Illumina and AVITI libraries to the reference genome, the duplication rate was calculated using Piccard with the MarkDuplicates command. Across all libraries, as the input DNA concentration decreased and the associated number of PCR cycles increased, the duplication rate also increased (Table 2). The 1/5th libraries had the highest rate of duplication for most libraries. We also observed larger variation in duplication rate across the AVITI libraries than was observed in the Illumina libraries. The full AVITI libraries often had lower rates of duplication than the Illumina libraries; as did the 4/5th libraries in some cases. Once the AVITI library prep miniaturization threshold dipped below 3/5th, the duplication rate was equal to Illumina libraries in the two libraries prepared from silica dried tissue, but drastically higher in the libraries prepared from fresh tissue.

**Table 2.**
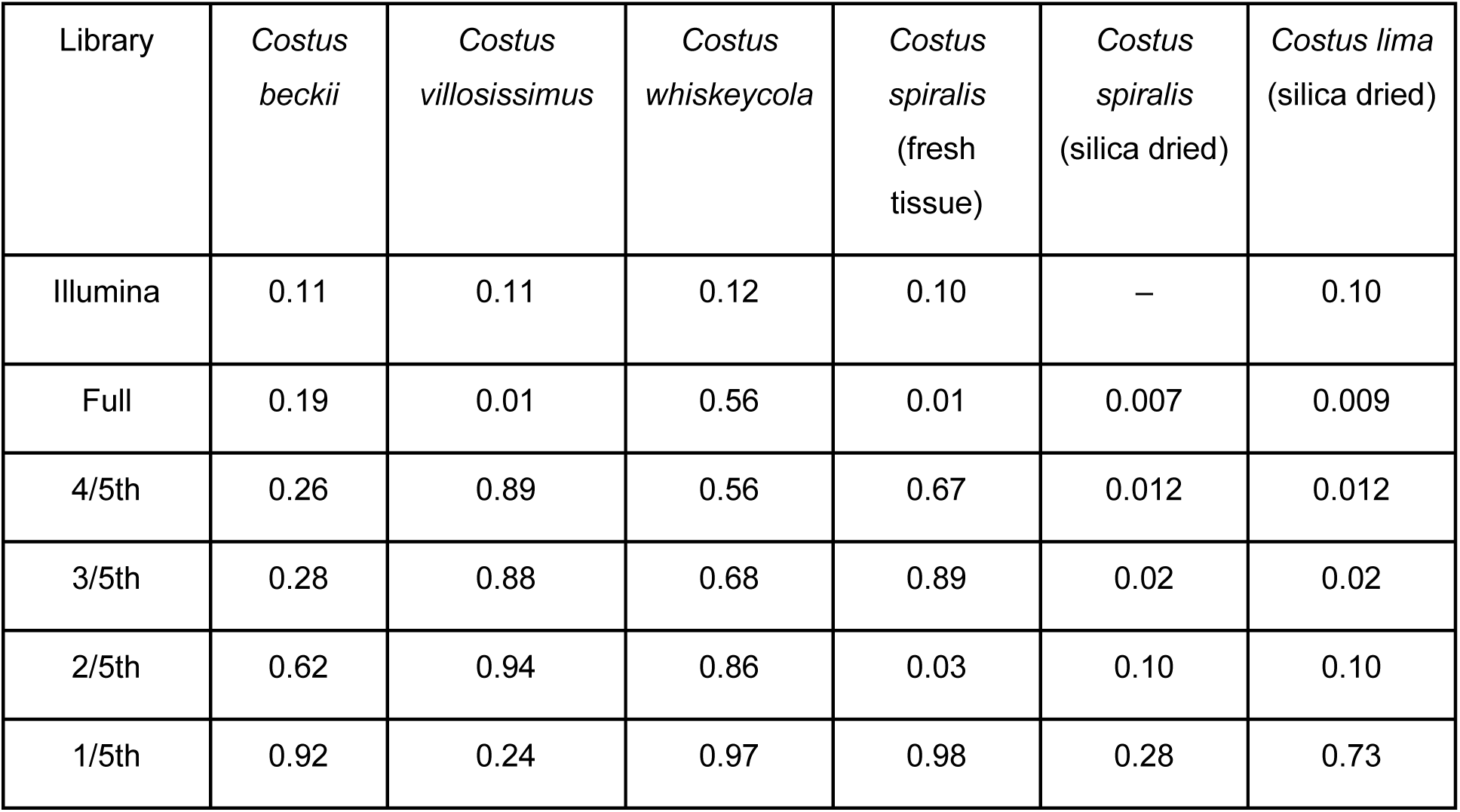
Duplication rate from each library as calculated by Piccard using the sorted BAM files after mapping cleaned reads to the reference genome.

| Library | *Costus beckii* | *Costus villosissimus* | *Costus whiskeycola* | *Costus spiralis*<br>(fresh tissue) | *Costus spiralis*<br>(silica dried) | *Costus lima*<br>(silica dried) |
| --- | --- | --- | --- | --- | --- | --- |
| Illumina | 0.11 | 0.11 | 0.12 | 0.10 | – | 0.10 |
| Full | 0.19 | 0.01 | 0.56 | 0.01 | 0.007 | 0.009 |
| 4/5th | 0.26 | 0.89 | 0.56 | 0.67 | 0.012 | 0.012 |
| 3/5th | 0.28 | 0.88 | 0.68 | 0.89 | 0.02 | 0.02 |
| 2/5th | 0.62 | 0.94 | 0.86 | 0.03 | 0.10 | 0.10 |
| 1/5th | 0.92 | 0.24 | 0.97 | 0.98 | 0.28 | 0.73 |

The resulting BAM files were also used to calculate the proportion of heterozygous sites across the genome. In terms of observed heterozygosity, the full and 4/5th AVITI libraries are close to the values inferred from the Illumina libraries (Table 3), which were sequenced at a much higher depth and likely better reflect the true heterozygosity estimate. As the amount of input DNA goes down for library prep, the estimated heterozygosity decreases, especially for the libraries prepared from fresh tissue. The reason for this association is likely due to sequencing output not deep enough to infer all heterozygotes. However, the libraries prepared from silica dried material are generally closer to the expectation of the Illumina data throughout the different miniaturization threshold with more equal sequencing output across libraries, with no library producing less than 8.5 Gbp of sequencing data.

**Table 3.**
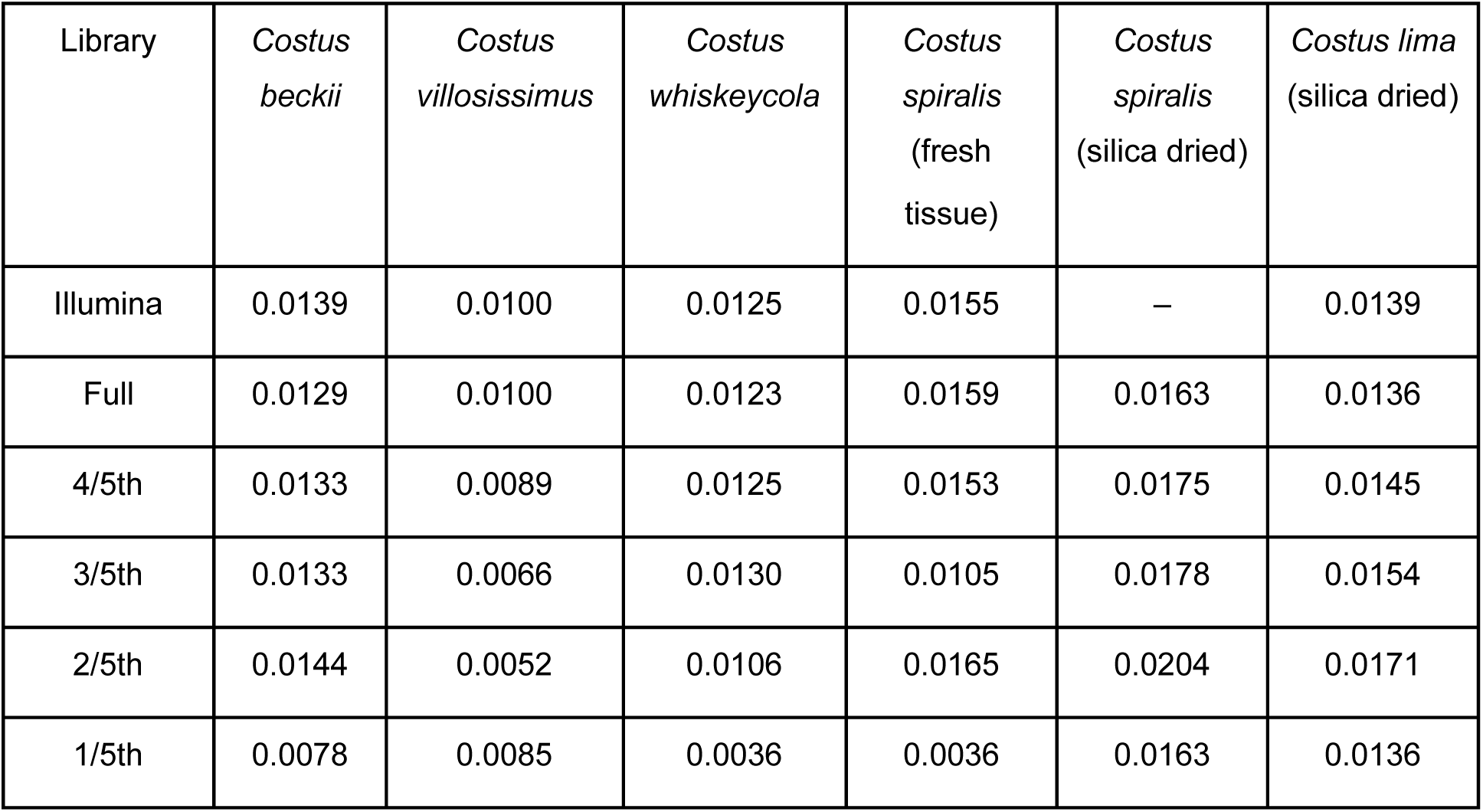
Proportion heterozygosity from all high-quality mapped sites as determined by ANGSD from the sorted BAM file after read mapping.

A total of 125,131,583 SNPs were initially called with BCFtools. After filtering to remove low quality SNPs with high amounts of missing data and LD filtering, a total of 639,116 SNPs remained for downstream analyses (Table 4). The Illumina libraries inferred little missing data, between 0.0029% and 0.08%; given that those libraries were sequenced close to 80x coverage, this was not unexpected. While the AVITI libraries exhibited higher levels of missing data, each of these were sequenced at much lower depth of coverage, ranging from 2.4 to 24.9 Gbp. As the input DNA threshold decreased, the depth of coverage for each sequencing library tended to be insufficient, resulting in an increase in the rate of missing SNPs after filtering. Most of the 1/5th libraries for the libraries prepared from fresh tissue had missing rates of over 92%, while the rate for the libraries generated from silica tissue was much lower (0.2 - 16.6%). Plotting sequencing output (Gbp) to the proportion of missing SNPs shows that once the total sequencing output achieved 10x coverage (∼10 Gbp), the amount missing SNPs was extremely low (Figure S2). The high level of missing SNPs is not just attributable to low sequencing coverage, since many libraries with low coverage did not have high levels of missing data. The libraries with the highest fraction of missing SNPs were *C. whiskeycola* 1/5th (0.9816), the fresh tissue library of *C. spiralis* 1/5th (0.9770), and *C. beckii* 1/5th (0.9292). The remaining five libraries with more than 50% missing SNPs were *C. villosissimus* 2/5th (0.9067), *C. villosissimus* 4/5th (0.8162), *C. villosissimus* 3/5th (0.8041), *C. whiskeycola* 2/5th (0.7088), and the fresh tissue *C. spiralis* 3/5th (0.6014).

**Table 4.** The fraction of missing SNPs after stringent filtering (--max-missing 0.8 --remove-indels --min-alleles 2 --max-alleles 2 --minQ 20 --minDP 3 --maf 0.05 --mac 3).

| Library | *Costus beckii* | *Costus villosissimus* | *Costus whiskeycola* | *Costus spiralis* (fresh tissue) | *Costus spiralis* (silica dried) | *Costus lima* (silica dried) |
| --- | --- | --- | --- | --- | --- | --- |
| Illumina | 0.0002 | 0.0003 | 0.0008 | 2.97e-05 | — | 0.0008 |
| Full | 0.0353 | 0.0022 | 0.0775 | 0.0002 | 0.0015 | 0.0158 |
| 4/5th | 0.0526 | 0.8162 | 0.1452 | 0.1543 | 0.0002 | 0.0094 |
| 3/5th | 0.0454 | 0.8041 | 0.2935 | 0.6014 | 0.0001 | 0.0042 |
| 2/5th | 0.2463 | 0.9067 | 0.7088 | 0.0001 | 0.0004 | 0.0073 |
| 1/5th | 0.9292 | 0.0168 | 0.9816 | 0.9769 | 0.0021 | 0.1664 |

**Figure S2.**
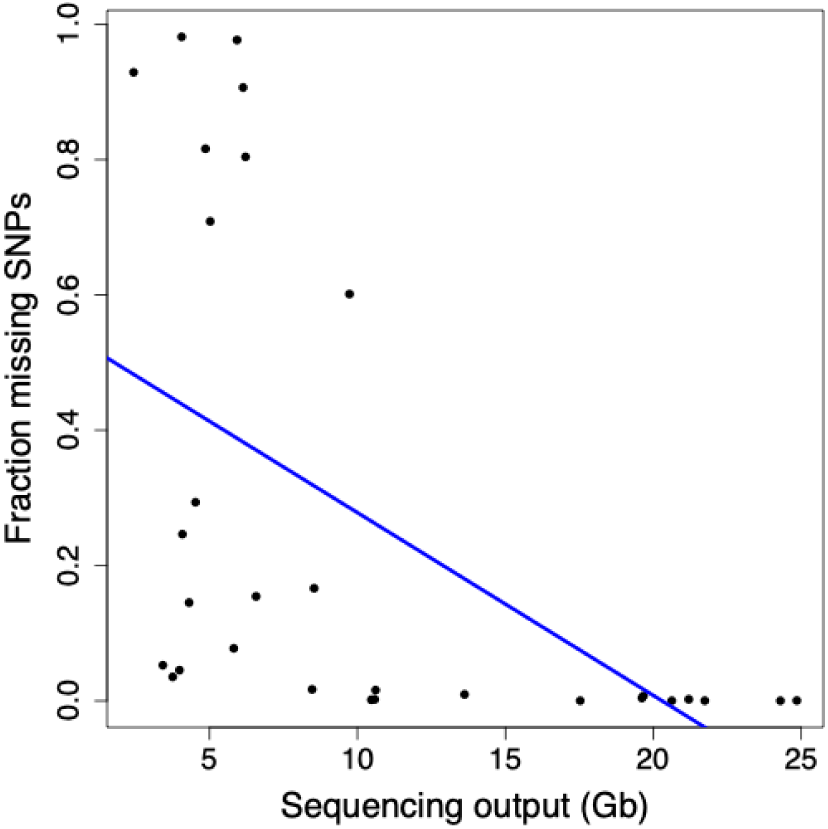
Regression analysis between sequencing output per library in terms of gigabase pairs versus the fraction of missing SNPs. The adjusted R2 value is 0.26.

After filtering SNPs so that only those present in a given accession were retained in the VCF file, Venn diagrams were inferred to determine the overlap across the five different AVITI libraries, as well as between the AVITI full and the Illumina libraries. For the *C. spiralis* fresh tissue libraries, only 6.3% (18,324 out of 290,633 SNPs) were found in all five libraries (Figure 3A). Seventy-three percent were found in everything but the 1/5th libraries (Figure 3A). Very few SNPs were unique to any given library with two unique SNPs found in the full library and ten found in the 2/5th libraries. Comparing the full library to the Illumina library, almost all of the SNPs were shared between the two (290,579 out of 290,633), with only 32 SNPs found just in the Illumina library and 22 found in just the AVITI full library (Figure 3B). For the *C. spiralis* silica libraries, the overlap between libraries was more even with 99.8% (290,136 out of 290,633 SNPs) shared between all five libraries; the 1/5th library for the *C. spiralis* silica tissue library was more representative of the other libraries with only 0.06% (184 out of 290,633 SNPs) shared across the other four libraries but missing in the 1/5th library (Figure 3C). Since we had both fresh and silica libraries from the same accession of *C. spiralis*, we were able to compare the full libraries of both to the Illumina library. Across the three libraries, all of the SNPs were accounted for in at least two libraries with no library having unique SNPs (Figure 3D). Twenty-two SNPs were missing in the fresh library, 225 SNPs from the silica library, and only 32 were missing from the Illumina library.

**Figure 3.**
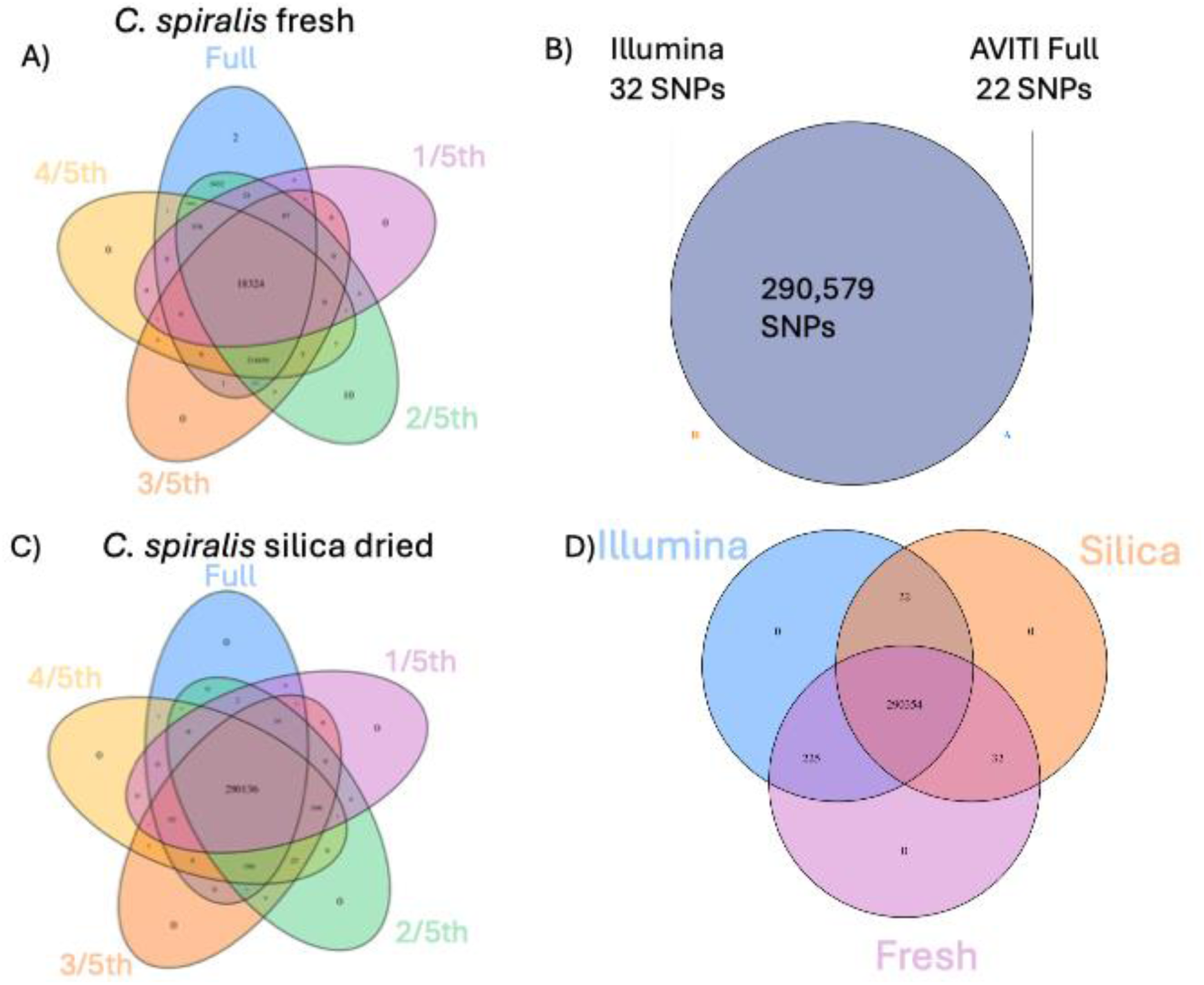
Venn diagrams showing shared versus unique SNPs across different libraries, with SNPs filtered to remove any site with missing data in a given library. Panel A shows the five AVITI libraries for the fresh *C. spiralis* libraries. Panel B shows the overlap in the Full AVITI library and the Illumina library for the fresh *C. spiralis* libraries. Panel C shows the overlap of the five libraries for the silica *C. spiralis*. Panel D shows the overlap of the Illumina, fresh *C. spiralis* AVITI full, and the silica *C. spiralis* AVITI full.

Overall patterns were slightly different across the remaining species when comparing just the AVITI libraries, especially when looking at libraries made with fresh samples as compared to silica samples, with very few SNPs unique to any given library (Figure S3). Across the fresh libraries of *C. beckii* and *C. whiskeycola*, 18.7% (54,318 out of 290,580 SNPs) and 7.2% (13,647 out of 290,465) were shared across all five libraries; with 71.9% (209,062 out of 290,580 SNPs) and 59.2% (171,939 out of 290,465) shared across all libraries except the 1/5th libraries, respectively (Figure S3A and C). The *C. villosissimus* libraries across the board had high rates of missing data in the 4/5th, 3/5th, and 2/5th libraries (Table 4), resulting in 17.1% (49,833 out of 290,619 SNPs) shared across all five libraries and 36.3% (105,312 out of 290,619 SNPs) shared between only full and 1/5th libraries (Figure S3B). The silica libraries of *C. lima* were similar to those of *C. spiralis,* with 90% (261,315 out of 290,497 SNPs) shared across all five libraries, with an additional 7.8% (22,612 out of 290,497 SNPs) missing in the 1/5th library which are shared by all others (Figure S3D).

**Figure S3.**
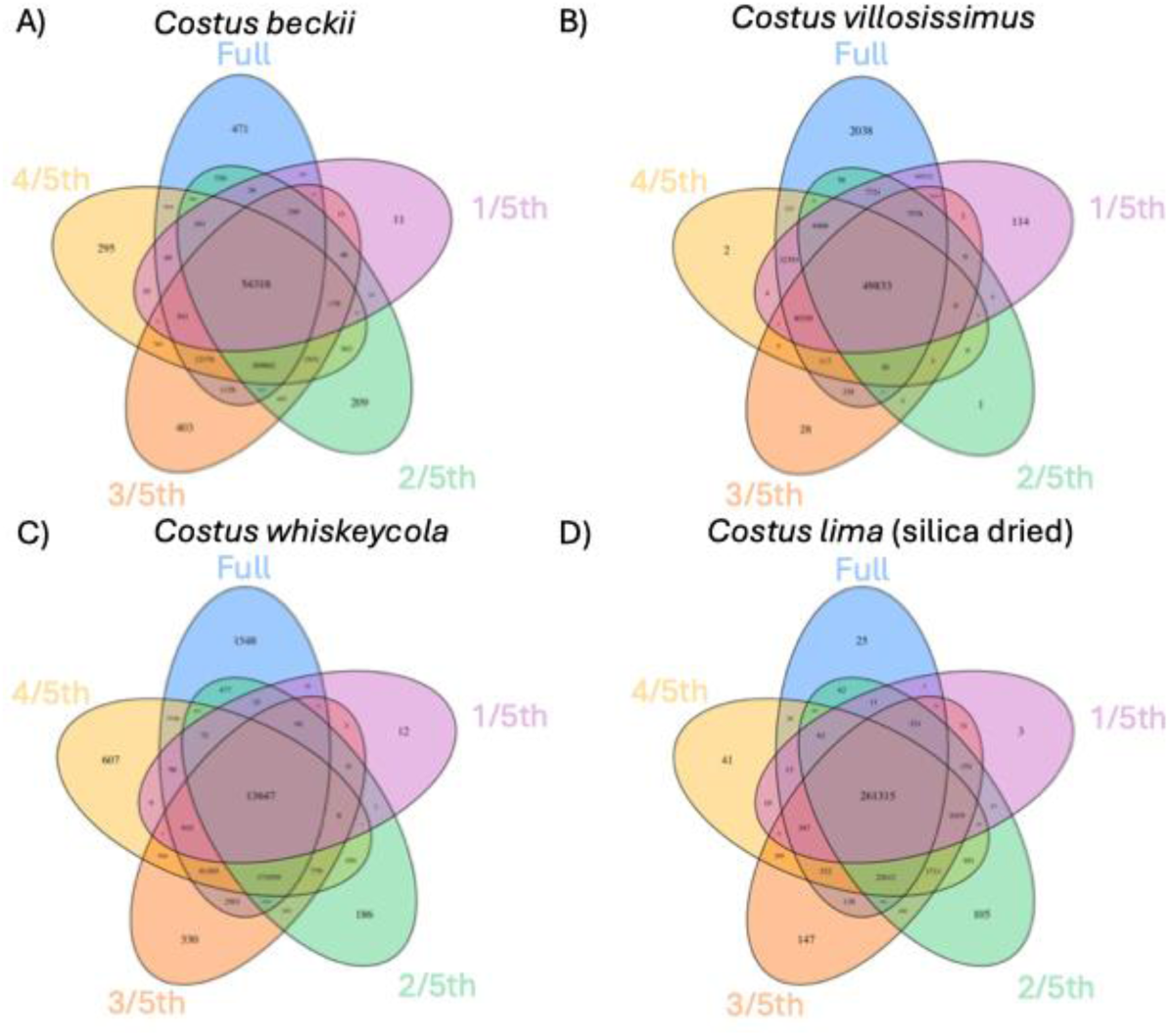
Venn diagrams showing shared versus unique SNPs across different libraries, with SNPs filtered to remove any site with missing data in a given library. Panel A shows the five libraries of *C. beckii*, Panel B shows the five libraries of *C. villosissimus*, Panel C shows the five libraries of *C. whiskeycola*, and Panel D shows the five libraries of silica *C. lima*.

When comparing SNP overlap between the Illumina and full AVITI libraries, the vast majority are shared across both. The biggest differences were found in *C. beckii* and *C. whiskeycola* for which the Illumina libraries had 5,335 and 5,330 unique SNPs, respectively, compared to 36 and 14 found in the AVITI data. *Costus villosissimus* had 302 unique SNPs in the Illumina library while the AVITI full library had 27; and *C. lima* was intermediate of the rest with 4,135 SNPs unique to the Illumina library and 104 unique to the AVITI full.

Even though the number of shared SNPs between most AVITI libraries, and specifically between Illumina and AVITI, are mostly overlapping, the SNPs in the different libraries are not identical, at least in terms of pairwise distance between samples (Table 5). Across all samples, the full libraries had a pairwise distance ranging from 1.1-2% compared to their Illumina counterparts. For most of the species, as the level of library miniaturization increased, so did the level of distance between that library and the Illumina library. The two silica libraries which showed the most consistent sequencing showed limited distance across all libraries, ranging from 1.9-2.5% in *C. spiralis* and 1.6%-2.2% in *C. lima*.

**Table 5.**
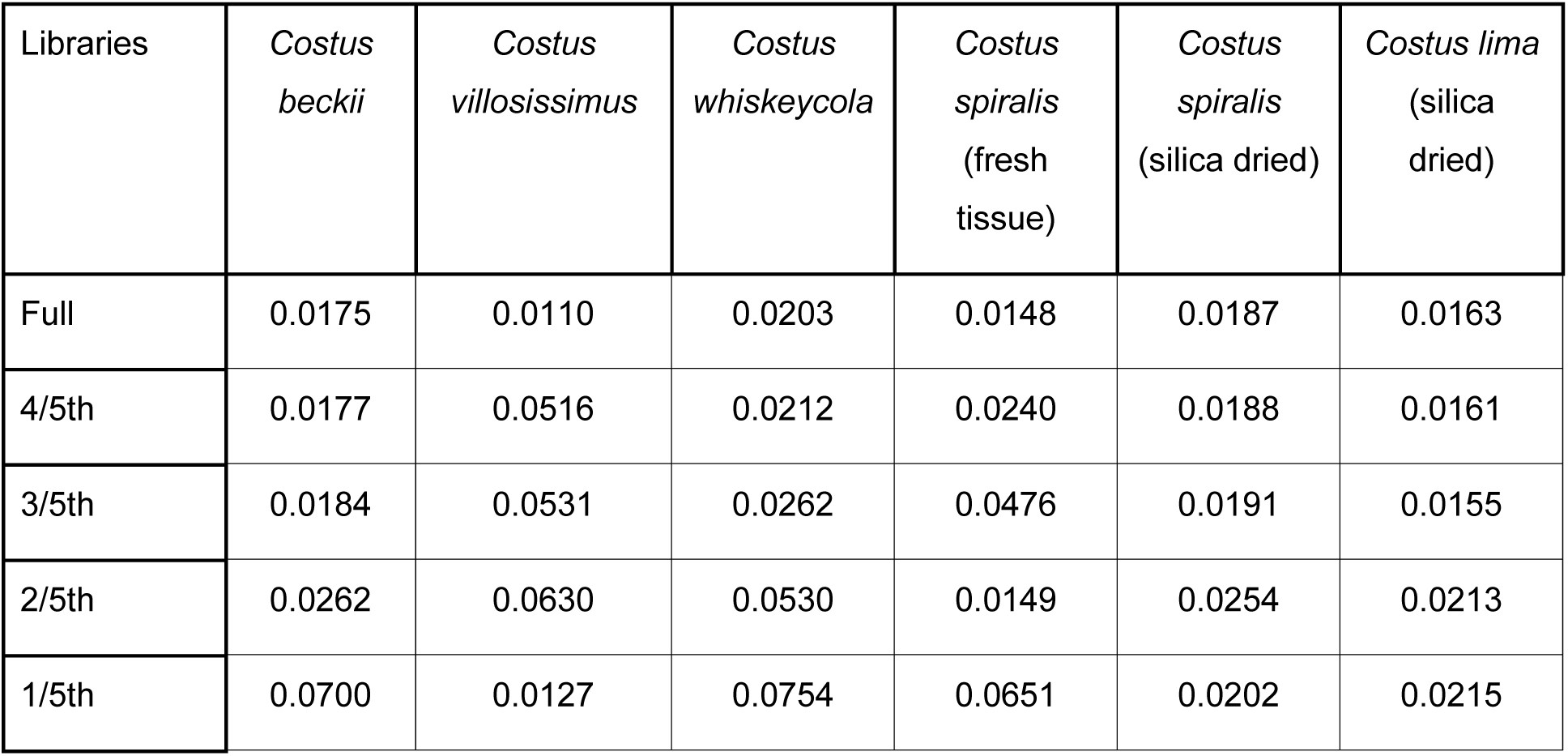
Pairwise distances of AVITI libraries compared to their Illumina counterparts as calculated from the FASTA conversion of the filtered SNP data set.

| Libraries | *Costus beckii* | *Costus villosissimus* | *Costus whiskeycola* | *Costus spiralis* (fresh tissue) | *Costus spiralis* (silica dried) | *Costus lima* (silica dried) |
| --- | --- | --- | --- | --- | --- | --- |
| Full | 0.0175 | 0.0110 | 0.0203 | 0.0148 | 0.0187 | 0.0163 |
| 4/5th | 0.0177 | 0.0516 | 0.0212 | 0.0240 | 0.0188 | 0.0161 |
| 3/5th | 0.0184 | 0.0531 | 0.0262 | 0.0476 | 0.0191 | 0.0155 |
| 2/5th | 0.0262 | 0.0630 | 0.0530 | 0.0149 | 0.0254 | 0.0213 |
| 1/5th | 0.0700 | 0.0127 | 0.0754 | 0.0651 | 0.0202 | 0.0215 |

### Downstream population genomics and phylogenomic analyses

The PCA generally showed good clustering of all libraries for the same accession, with the outliers tending to be the 1/5th libraries across the different principal components (PCs)(Figure 4). The four top PC contributors explained the following amount of variation: 20.71% (PC1), 12.94% (PC2),10.46% (PC3), and 6.98% (PC4). The Illumina libraries (open circles) tend to be positioned closest to the AVITI full (filled circles), AVITI 4/5th (square), or AVITI 3/5th (triangle) for all samples except for *C. villosissimus*, which suffers from high amounts of missing data (Table 4).

**Figure 4.**
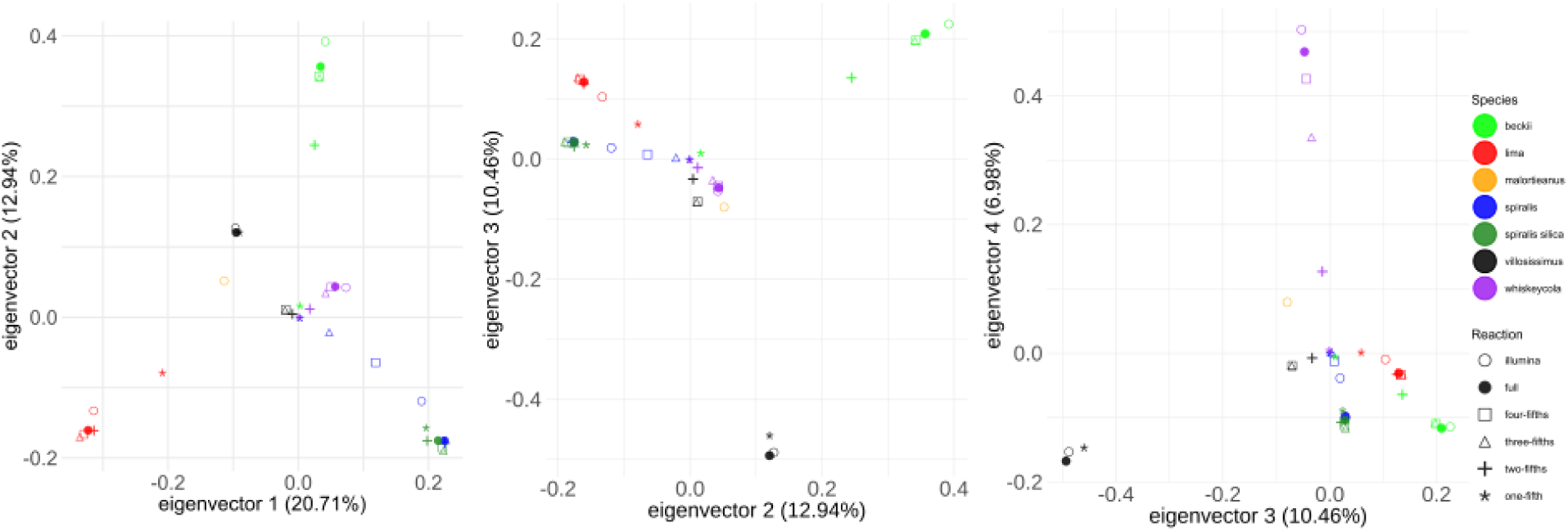
Principal component analysis with the filtered SNP setting including the Illumina library and the five AVITI libraries for each accession, as well as *C. malortieanus* which serves as the outgroup for other analyses. The three different panels represent PC1 vs PC2, PC2 vs PC3, and PC3 vs PC4.

The Structure analyses performed In LEA showed that the best K value is either K=5 or K=6 based on cross-entropy (Figure S4). Structure barplots showing the level of admixture for each library, with high degrees of similarity in most libraries per accession, except for the AVITI 1/5th libraries (Figure 5). Across all libraries, the AVITI Full libraries are most similar to the Illumina libraries than any of the other scaled AVITI libraries.

**Figure 5.**
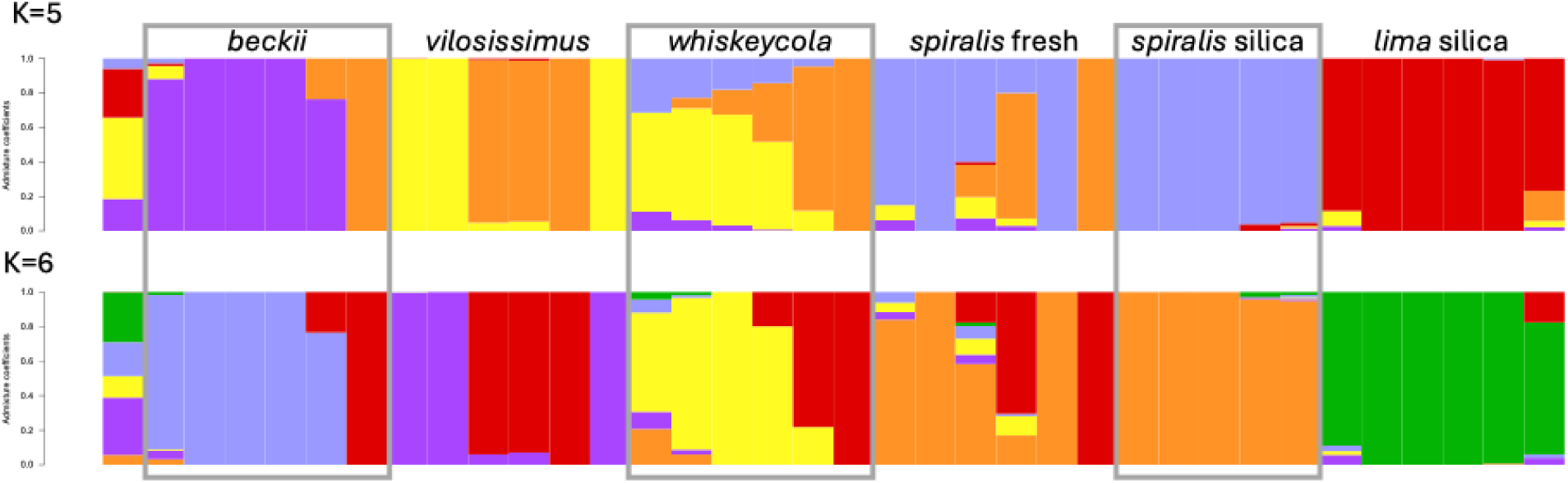
Structure barplots produced by LEA showing K=5 and K=6. The six libraries for each of the five accessions are shown by boxes. Across most accessions, the AVITI 1/5th libraries are assigned to different ancestral groups. The order of libraries for accessions are: Illumina, Full, 4/5th, 3/5th, 2/5th, and 1/5th, except for the silica *C. spiralis* libraries that do not have the Illumina library grouped and the outgroup *C. malortieanus*.

**Supplemental Figure S4.**
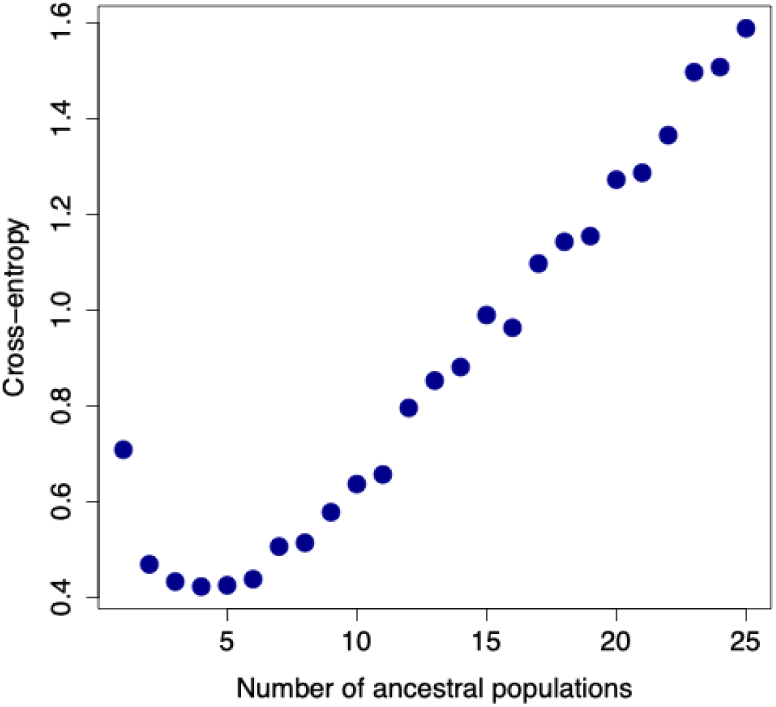
The cross-entropy plot for the best K in LEA, showing K=5 and K=6 being highly similar.

The RAxML phylogenetic analyses using 639,116 SNPs showed strong clustering of all libraries from the same accession, regardless if the libraries were Illumina or AVITI (Figure 6). In all cases, all six libraries (Illumina plus the five AVITI libraries) produced monophyletic groups with 100% support; except for the case of the fresh and silica *C. spiralis* libraries that were interspersed. However, both the fresh and silica libraries of *C. spiralis* came from the exact same accession. Pruning the sampling down to just the taxa with both sets of libraries showed the same basic topology, with strong support at most nodes (Supplemental Figure S5). Only including the Illumina and AVITI Full libraries showed 100% bootstrap support for sister relationships for all targeted taxa, except for *C. spiralis* where the two AVITI libraries were sister to each other, and the Illumina library was subsequently sister to this clade (Supplemental Figure S6).

**Figure 6.**
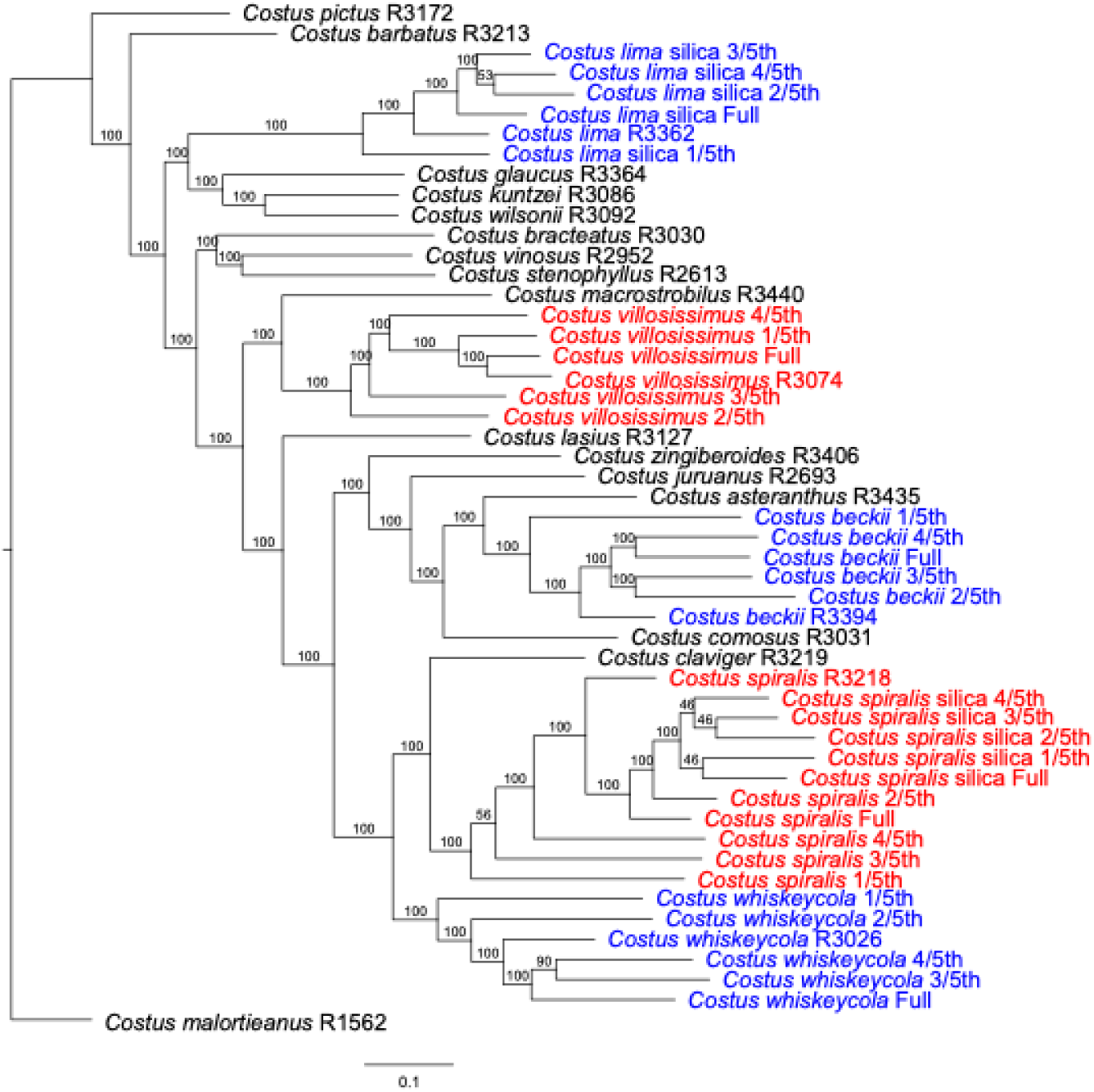
RAxML phylogenetic tree using the GT model of molecular evolution with all *Costus* accessions showing strong bootstrap support for clustering of all libraries from the same accession, regardless if they were Illumina or different AVITI thresholds.

**Supplemental Figure S5.**
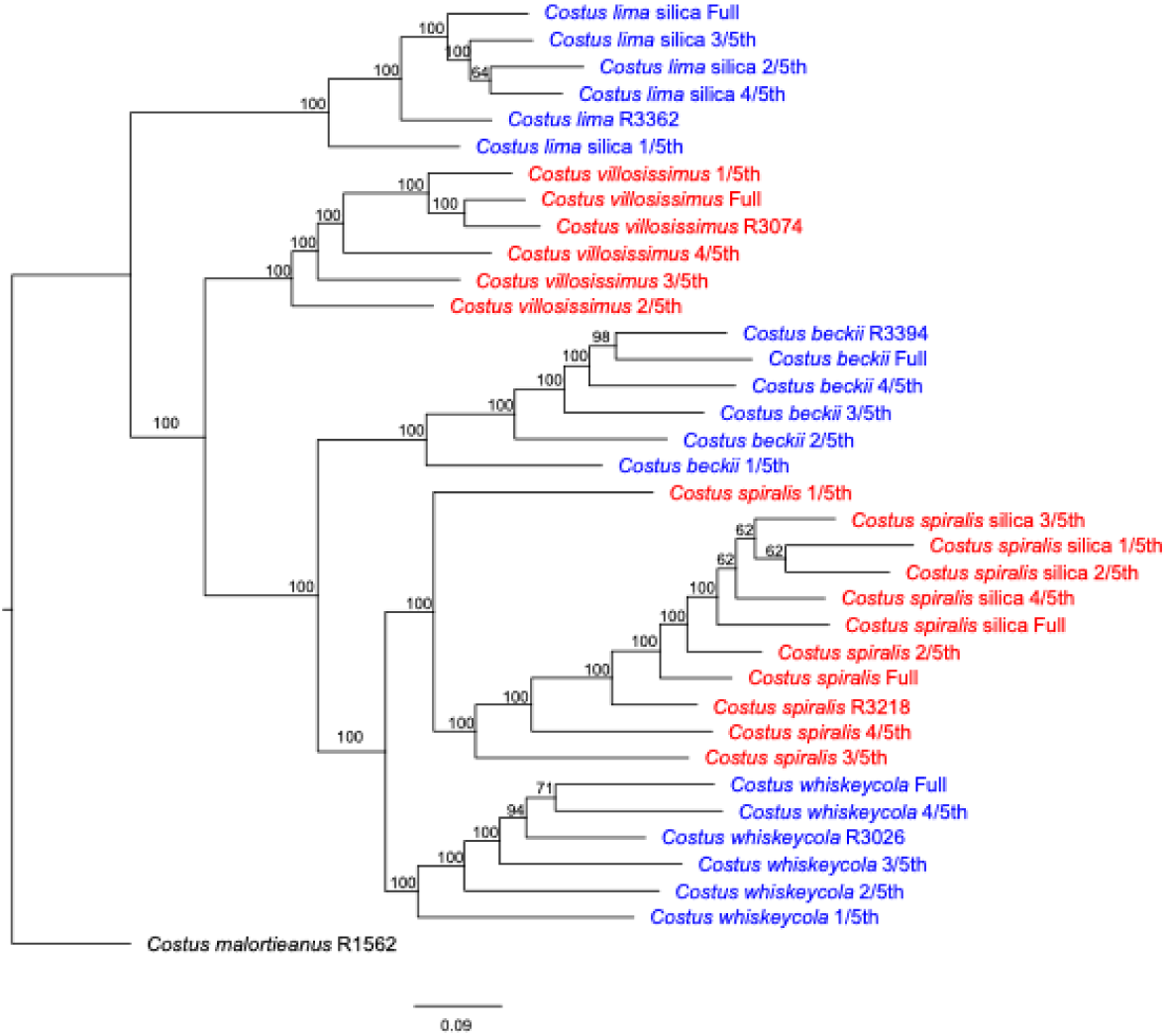
Pruned phylogenetic analyses in RAxML with only the taxa that included both Illumina and AVITI libraries, with the exception of *C. malortieanus* (Illumina only) that was used as the outgroup.

**Supplemental Figure S6.**
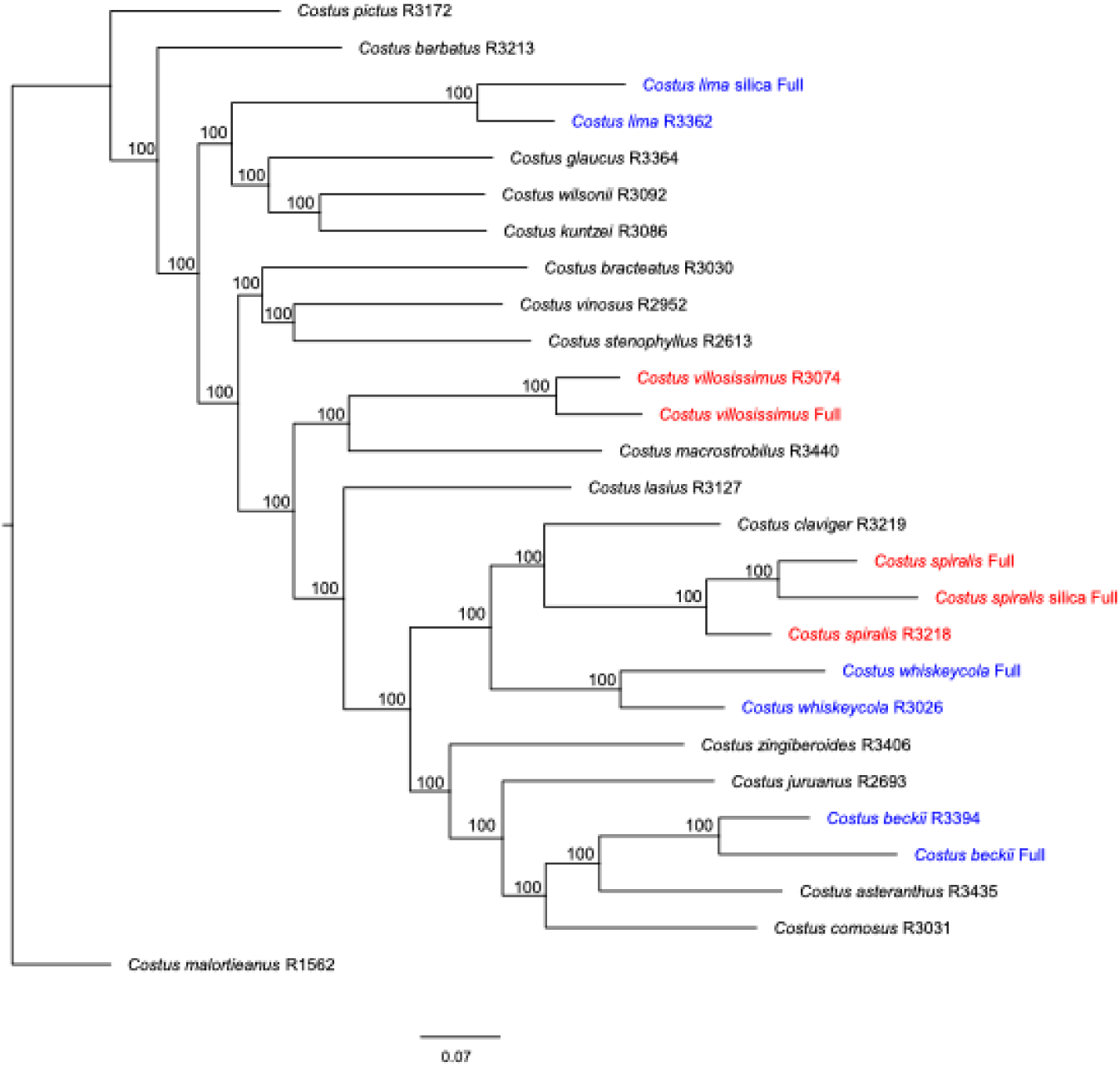
Pruned phylogenetic analyses in RAxML with only the Illumina and AVITI full libraries for the targeted accessions to ensure that phylogenetic clustering was not occurring based on the denser sampling of certain taxa.

## Discussion

This study set out to address two primary questions regarding the use of AVITI sequencing: can the data be paired with Illumina data to infer biologically meaningful interpretations and what are the impacts of library miniaturization on sequencing reads, SNP inference, and downstream analyses. We showed that as library miniaturization was reduced to smaller volumes, the number of PCR cycles needed for library amplification went up, directly resulting in the increased read duplication. This increased level of duplication also contributed to an increased level of missing SNPs across libraries. The level of observed heterozygosity across the genome was mostly consistent across libraries of the same accession, with an apparent underestimation in the smaller library volumes partially attributed to the increased duplication rate. Similarly, the number of SNPs shared across libraries of the same accession were mostly consistent across libraries, except in cases where the 1/5th libraries exhibited the most missing data. Phylogenomic analyses showed strong agreement between Illumina and AVITI libraries, even at the lower miniaturization volumes. Clustering with PCA and Structure were mostly consistent across all libraries, with the inclusion of the 1/5th libraries being the most varied.

When comparing SNP retention across platforms and across miniaturization thresholds, we found strong concordance in the SNPs that were retained. Across miniaturization levels, the majority of SNPs were shared across all five, or in some cases four of the five, levels (Figure 3 and Figure S3), with very few SNPs unique to a single stage. The most drastic instances of high levels of unique SNPs were in the full library volume of *C. villosissimus* having 2,038 unique SNPs compared to the other AVITI libraries and in *C. whiskeycola* where the full AVITI library had 1,548 unique SNPs compared to the other miniaturization thresholds. In both these instances, the full libraries of both species had drastically less missing data than most of the other libraries for that species (Table 4). The SNP retention between the full AVITI library and the Illumina library were almost entirely identical (Figure 3B), with only 0.019% of SNPs unique to either the AVITI or the Illumina library in *C. spiralis*. The percentage of shared SNPs between platforms was much higher than previously shown by de Ronne et al. [5], who showed 62.4% overlap in SNPs, with Illumina harboring about one third more unique SNPs than were unique to AVITI. However, the lack of shared SNPs in their study was attributed to the use of GBS approach instead of a whole genome sequencing approach, which has a higher inherent variability of the sites/loci sampled, in addition to variation in sequencing depth and sequencing chemistry between platforms [5]. Differences in retained SNPs between platforms is also impacted by sequencing quality, especially in tandem repeats and homopolymers, in which cases AVITI has a higher accuracy than Illumina [4].

Miniaturization during library preparation is a commonly employed method for cost savings in NGS experiments, in terms of time, amount of plastics used, and needed volume of reagents [14,46]. Pillay et al. [11] found a 98% sequencing success rate when using 1/10th reactions, however in our smallest miniaturization of 1/5th reactions we found an exceptionally high level of duplication (92-98%) in several of the libraries (Table 2) accompanied with a high rate of missing SNPs (92.9-98.2%) after stringent filtering (Table 4). These levels are in strong contrast to the full volume libraries for the same accessions that showed duplication rates from 1-56% and percentage of missing SNPs ranging from 0.22-7.8%. The high levels of read duplication are similar to a recent study using RNA-Seq instead of whole genome sequencing which showed a strong negative correlation with input concentrations less than 125 ng and the observed duplication, with up to 96% reads discarded as duplicates [47]. The different miniaturization levels were carried out to keep the reaction volumes consistent, however an alternative and likely better approach is to keep the concentrations of enzymes the same and reduce volumes. Reduction to small volumes, especially at the 1/5th volume, can lead to issues due to pipetting errors and viscosity of reagents [48]. Additionally, miniaturization attempts that do not require enzymatic fragmentation are likely easier than the Elemental Elevate kit used here. The fragmentation step was critical in the success of library construction; if the fragmentation reaction was not mixed sufficiently well by pipetting up and down 10 times, we saw a total failure in the reduction of DNA size and libraries with little to no qubit readings. In fact, even libraries at the full volume did not always fragment appropriately, with poorly performing libraries in terms of sequencing output showing less uniformity in final library size than those that generated 10+ Gbp of data (Supplemental Figure S1).

Even with some of the technical difficulties in the more extreme miniaturization levels tested here, many of the analyses showed strong congruence in both SNPs called and downstream analyses. Across the worst libraries of *C. beckii* and *C. whiskeycola*, only 18.7% and 7.2% of SNPs were shared across all five libraries, respectively; with 71.9% and 59.2% shared across all libraries except the 1/5th libraries (Figure S3A and C). Better producing libraries showed over 90% overlap in SNPs across all five AVITI libraries. The proportion of observed heterozygosity was mostly consistent across all libraries within a species (Table 3), with a reduced level observed in the 1/5th libraries most likely attributed to the higher duplication (Table 2) due to the increased number of PCR cycles during library amplification (Table 1). The 1/5th miniaturized libraries were resolved in the same clade as the other samples from the species, but were often observed to be the first diverging member in that clade (Figure 6), which is also supported by the higher pairwise distance between the 1/5th libraries and the Illumina libraries (Table 5). The Structure plots also displayed a tendency to place the 1/5th libraries in a different ancestral group compared to the other library volumes, as observed in *C. beckii, C. spiralis*, and *C. whiskeycola*, or with a higher level of admixture as observed in the silica samples of *C. lima* (Figure 5). The extent that these patterns are upheld across different library kits or sequencing platforms is largely unknown since results from population genomic and phylogenomic analyses are not typically shown in studies comparing library miniaturization.

Using the AVITI for genome sequencing should be seen as a viable option for phylogenomic and population genomic analyses, even when previous Illumina data exists and the goal is to enlarge the dataset with additional taxon sampling. We see strong agreement in both the inferred SNPs across libraries but also in inferred evolutionary relationships using a multitude of commonly applied analysis methods. While library miniaturization can be seen as a cost cutting measure, there are drawbacks when scaling down too much including increased read duplication, increased levels of missing data, and potentially misleading results in terms of heterozygosity or evolutionary relationships.

## Data availability

Raw Illumina reads are available from the Sequence Read Archive with the SRR numbers SRR18516534-SRR18516558. AVITI fastq files are available with SRR numbers XX-XX. The *Costus bracteatus* genome is available from CoGe with ID 63651.

## Author contributions

DA, EH, JBL, JJH, and JMF generated the data; JBL and JMF analyzed the data; JBL and CDS designed the experiment; JBL wrote the first version of the manuscript; all authors revised and approved the final version.

## Acknowledgements

The authors thank Timm Hamp for helping secure sequencing reagents and troubleshooting protocols, without him the project would not have been possible; as well as Joel Garcia and other support staff at Element Biosciences for answering questions about library prep and demultiplexing. The authors also thank Amy Lyndaker and William Lai for assistance running the AVITI sequencing runs. The concept of the project and results emerged from a graduate seminar Special Topics in Plant Sciences PLSCI 6940 offered by JBL, supported by Cornell Provost funds for teaching provided to CDS.

